# Chenonceau: mesoscopic MRI deep phenotyping of a post-mortem human brain at 7 and 11.7 Tesla

**DOI:** 10.64898/2026.05.26.726838

**Authors:** S. Legeay, J. Beaujoin, R. Yebga Hot, A. Popov, I. Uszynski, B. Herlin, F. Poupon, I. L. Maldonado, C. Destrieux, C. Poupon

## Abstract

The human brain’s organization, from macroscopic connectivity to microscopic architecture, has never been mapped across multiple modalities in a single post-mortem specimen at mesoscopic resolution. Here we present Chenonceau, a complete post-mortem human brain atlas acquired using two ultra-high-field preclinical MRI systems. Combining anatomical, diffusion, and quantitative MRI from 100 to 200 μm isotropic resolution, the 8000-hour acquisition campaign overcomes the fundamental trade-off between spatial resolution, tissue coverage, and multimodality that commonly constrains MRI neuroimaging in humans. The resulting open-access dataset integrates the first whole-brain mesoscopic connectome along with cortical lamination, myelo-, and cyto-architecture mapping. Importantly, the spatial and angular resolutions achieved reveal the intracortical connectivity at the whole-brain scale. Bridging the scale gap between in vivo imaging and post-mortem microscopic datasets, the Chenonceau brain atlas establishes a foundational reference for joint mesoscale analysis of human brain structure, connectivity, and microarchitecture. The dataset is openly available on the EBRAINS portal.

## Main

Fine mapping the microarchitecture of a complete post-mortem human brain, including its myeloarchitecture, cytoarchitecture, and nested connectome, remains a major challenge. From the seminal atlases produced by anatomists in the last century, through dissections and histology (*1–4*), to the latest technological innovations in ultra-high-field magnetic resonance imaging (UHF-MRI), optical, X-ray, electronic and immunohistochemistry-based (IHC) microscopy (*5–8*), and mass spectroscopy (*9*), it is now possible to cover a broad range of scales and contrasts essential for a global understanding of the brain structural organization. Although microscopic approaches appear closer to the ground truth, they require considerable investment to scan and reconstruct the entire brain (*10*). Preparatory steps, such as sectioning, IHC, or staining, often remain obstacles to multimodal imaging.

Benefiting from extended scan times, *post-mortem* UHF-MRI achieves mesoscopic resolution and offers the possibility of addressing multimodality, facilitating the exploration of gross anatomy and the concomitant characterization of microstructure. T1- and T2-weighted quantitative MRI (qMRI) provide insights into the myeloarchitecture (*11, 12*), while T2*-weighted qMRI gives access to magnetic susceptibility and therefore reveals the iron content of structures (*13, 14*). Finally, diffusion MRI (dMRI) has become a key tool for inferring human brain structural connectivity and characterizing the cytoarchitecture of brain structures by observing the anisotropy and time dependency of water movement in tissues (*15–22*).

Beyond multimodality, UHF-MRI can be combined with various microscopic exploration techniques, such as 3D-PLI, OCT, or Nissl stainings, to navigate across scales, as recently demonstrated in the macaque brain (*23*), which is tenfold smaller in volume than the human brain. In humans, recent whole-brain studies have shown the potential of UHF-MRI to reach the mesoscale for anatomical MRI (aMRI) and qMRI (*24, 25*) and the half-millimetre scale for dMRI (*26–30*). Unfortunately, the resolution of dMRI remains insufficient to map the nestedness of the human brain connectome (31) and the cytoarchitecture of its finest structures, such as cortical layers.

Leveraging higher magnetic fields alongside stronger gradients, small-animal MRIs push resolution beyond what is achievable on clinical MRIs today. While several studies have demonstrated this, focusing on the hippocampus, brainstem, and cerebral cortex (*32–37*), the small diameter of the magnet tunnel on these instruments, as well as the sensitivity profile of their antenna, restricts these studies to small samples. To cover the entire brain, it is necessary to cut it into blocks to run a large-scale acquisition campaign by scanning each block individually, and to develop a pipeline to recompose the whole brain from its blocks. This severe limitation is the price for a dramatic improvement in the signal-to-noise (SNR) and contrast-to-noise ratios. Because each sample is much smaller than the B1 radiofrequency wavelength, it avoids the usual destructive dielectric interference observed at extreme fields, thus preserving contrast. Similarly, the configuration of preclinical antennas relative to the small brain samples ensures a maximum distance of 2 to 3 cm from the receiving coils, thereby enhancing the SNR.

The Chenonceau study presented here followed this “split-recomposition” strategy. This extensive deep phenotyping initiative focused on a single *post-mortem* human brain, named Chenonceau, after the geographical origin of its donor near the famous French château. The acquisition campaign, conducted on NeuroSpin’s preclinical 7T and 11.7T MRI scanners, lasted nearly 8,000 hours. The outstanding large-scale dataset provides, for the first time, multimodal UHF-MRI of an entire human brain at unprecedented resolutions of 100 μm for anatomical MRI and 200 μm for quantitative and diffusion MRI. The brain specimen was obtained from a 92-year-old man with a history of macular degeneration and mild cognitive disorders, enrolled in the Body Donation program of Tours University. The study was approved by the Local Ethical Committee (approval 2026-016) and declared to the French national CODECOH, which controls human tissue collections. After fixation, the brain was split into 13 parallelepipedal block samples that fitted within preclinal MRI antennas.

## Results

### Large-scale UHF-MRI acquisition campaign on the Chenonceau brain

Mesoscopic aMRI and dMRI datasets were collected using the NeuroSpin’s preclinical 11.7T MRI scanner (Bruker BioSpin, Ettlingen) equipped with a powerful gradient system (maximum amplitude of 760 mT.m^-1^ and slew rate of 9500 T.m^-1^.s^-1^) using a 60 mm in diameter volume birdcage antenna to ensure a homogeneous sensitivity over a rectangular field of view (FOV) of 42×42×56 mm^3^. The complete coverage of a block required three to four imaging sessions, depending on the length of the brain block along the rostro-caudal direction.

Each imaging session included two anatomical T2-weighted spin-echo MRI scans using 2D and 3D encoding schemes with respective isotropic resolutions of 150 and 100 μm, and 17 dMRI scans with an isotropic resolution of 200 μm using a segmented 3D Pulsed Gradient Spin Echo echoplanar (EPI) sequence. The 17 dMRI scans included 175 diffusion-weighted volumes, corresponding to a multiple-shell hybrid q-space sampling over three spheres at b = 1500/4500/8000 s.mm^-2^ along 25/60/90 diffusion directions uniformly distributed over the unit sphere (*38*). The complete acquisition of a single FOV required 107 hours, leading to a massive acquisition campaign totaling 4,815 hours over two years to gather the 45 aMRI and dMRI FOVs needed to cover the entire brain.

In parallel, quantitative T1-, T2-, and T2*-weighted MRI data were collected on the NeuroSpin’s preclinical 7 Tesla MRI scanner (Bruker BioSpin, Ettlingen) equipped with a 740 mT.m^-1^ gradient system (maximum slew-rate of 3440 T.m^-1^.s^-1^) and a 60 mm volume antenna. We used the 7 Tesla scanner instead of the 11.7 Tesla scanner to enable parallel acquisition campaigns on both imagers, thereby reducing the acquisition campaign duration. The same acquisition strategy was adopted for the 13 blocks, with each block being successively scanned at 11.7T and 7T.

The imaging session for each FOV comprised 1) a series of 65 T1-weighted MRI volumes acquired using a variable-flip-angle FLASH sequence (VFA-SPGR), 2) a series of 30 T2-weighted MRI volumes acquired using a multiple-spin-multiple-echo MSME sequence, 3) a series of 10 T2*-weighted MRI volumes acquired using a FLASH SPGR sequence, and 4) a B1+ radiofrequency field calibration scan acquired from a double flip angle 3D EPI spin echo sequence. All 105 quantitative MRI volumes were acquired at the exact 200 μm isotropic resolution used for dMRI data, with 3D Fourier-space encoding to maximize SNR.

Acquiring a single qMRI FOV required 66 hours, *i*.*e*., 3,168 hours for the entire Chenonceau brain. Particular care was taken to preserve the blocks by rehydrating them monthly throughout the 2-year acquisition campaign. A quality control process was established to ensure no drift in tissue or image quality (see supplementary text 1). Fig. 1 summarizes the sample preparation and both aMRI/dMRI and qMRI acquisition campaigns.

**Fig. 1:**
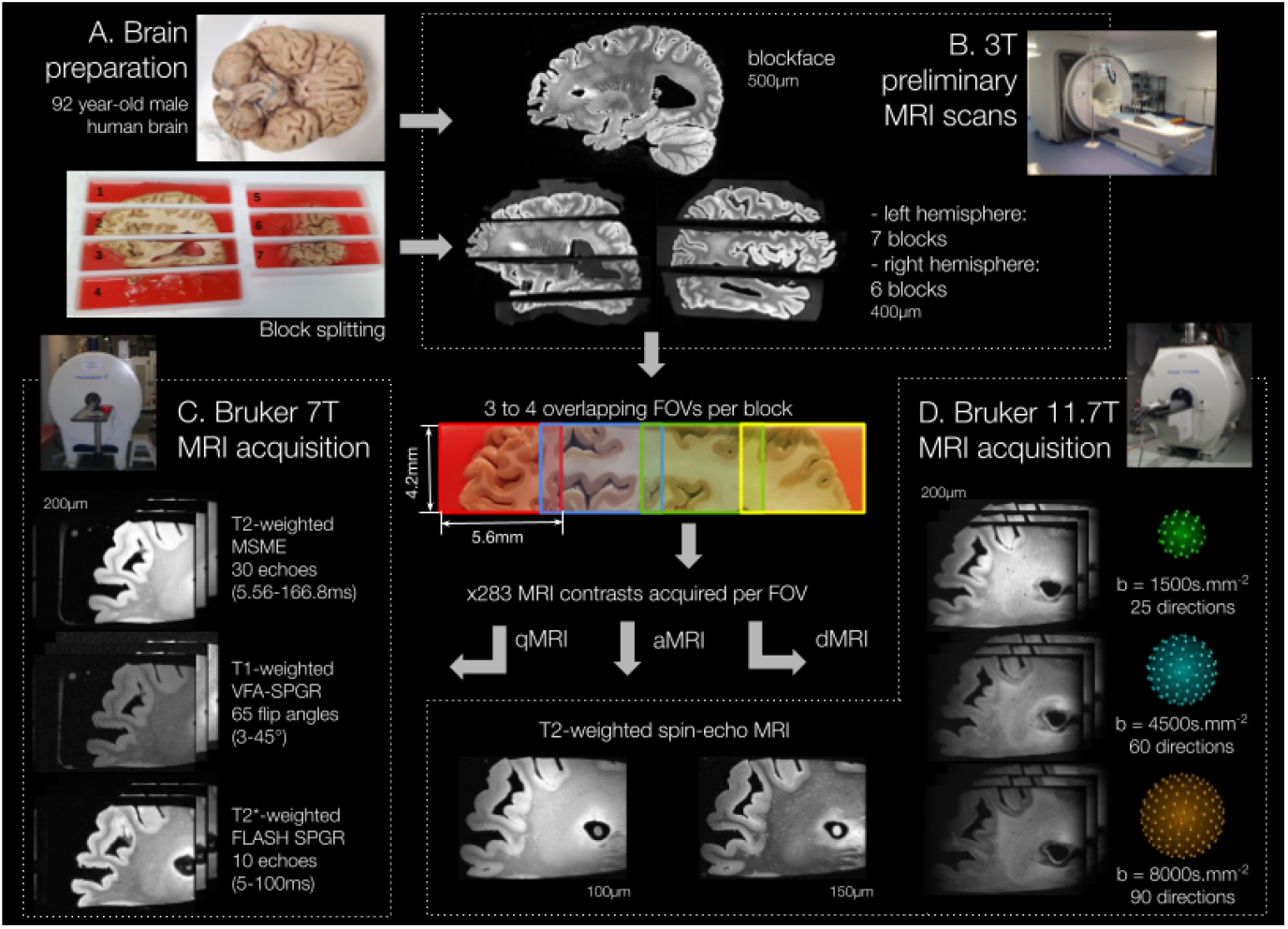
Brain sample preparation and MRI scanning campaigns at 7T and 11.7T. After scanning the whole brain at 3T to get an anatomical blockface volume, the two hemispheres of the Chenonceau brain were cut into 13 blocks. Intermediate images of the blocks were performed at 3T. Each block was split into 3 to 4 fields of view that were individually scanned on the two 7T and 11.7T MRI systems. The acquisition of a single FOV required a scan duration of 107 hours at 11.7 Tesla and 66 hours at 7 Tesla, totaling almost 8,000 hours over 2 years to gather the fifty or so FOVs needed to cover the entire brain.

**Fig. 2:**
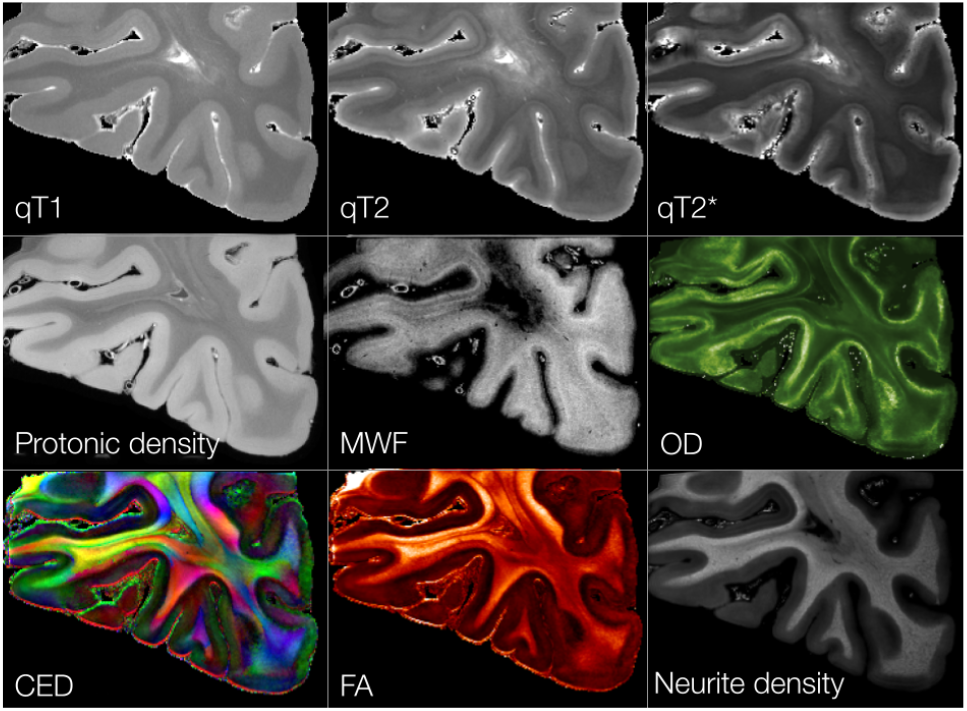
Myelo- and cyto-architectonic features extracted from qMRI and dMRI data in one FOV (occipital lobe of the right hemisphere). Voxel-wise microstructural features were derived from single-compartment models (qT1, qT2, qT2*, and corresponding protonic density) and from multi-compartment models of qMRI (myelin water fraction (MWF) (*12*)) and of dMRI (neurite orientation dispersion (OD) and neurite density indices stemming from the NODDI model (*20*)). Color-encoded (CED) and fractional anisotropy (FA) maps were derived from the DTI model.

### Myelo- and cyto-architectonic features extraction

T1, T2, and T2* maps were derived from their respective acquisition. Benefiting from a wide range of echo times in T2-weighted MRI and flip angles in T1-weighted MRI, the multi-compartment analysis of quantitative data provided valuable insight into the tissue myeloarchitecture. In the case of a *post-mortem* sample deprived of cerebrospinal fluid (CSF), three compartments were taken into consideration: the myelin-related compartment and two distinct gray and white matter compartments. Largely adapted from models in (*12*), the myelin water fraction (MWF) was robustly estimated in the entire brain. Fractional anisotropy, mean diffusivity, and color-encoded direction maps were computed using a diffusion tensor imaging (DTI) model. In addition, by leveraging the three diffusion shells, cytoarchitectonic markers, including neurite orientation dispersion and neurite volume fraction, were estimated using the NODDI model.

### Brain reconstruction: from pieces to puzzle

A total amount of 3.6 TB of raw data was collected for all aMRI, dMRI, and qMRI scans. The analysis of this unique multimodal MRI dataset required the development of a dedicated distributed image processing workflow (Fig. 3) to 1) correct the different imaging artifacts (geometric distortions, noise, intensity bias) of each FOV in all MRI modalities (see supplementary text 2), 2) compute the diffeomorphic transformation graph allowing the repositioning of all FOVs in a common space, corresponding to a total of 270 diffeomorphic registrations, and 3) finally recompose the target whole brain MRI dataset at the mesoscale for all modalities.

**Fig. 3:**
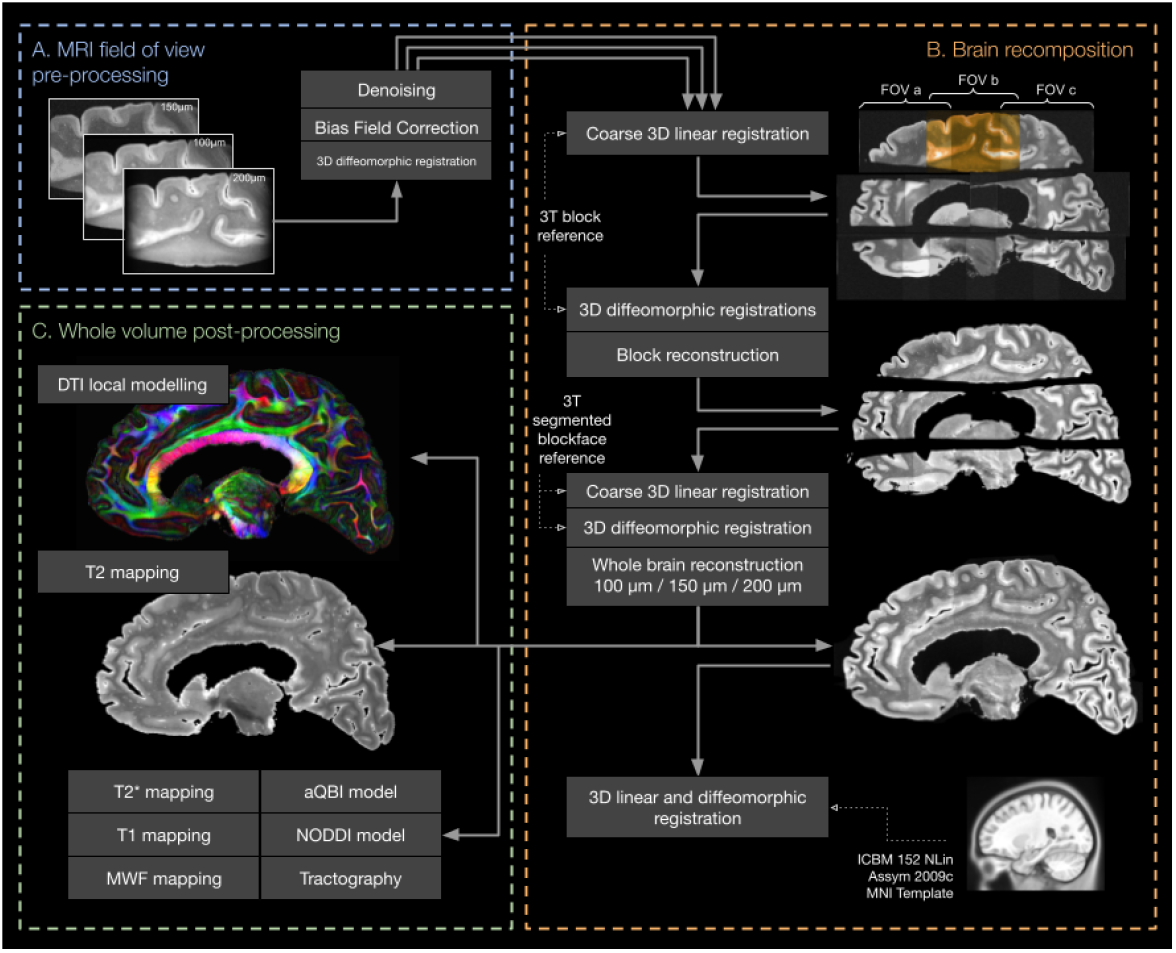
Processing of the Chenonceau brain from FOVs pre-processing to post-processing at the level of the entire brain. (A) Initial steps involved standard MRI artifact removal, including bias field correction and noise filtering (see supplementary text 2), followed by (B) the meticulous registration of the fields of view into the blockface space and the asymmetric non-linear ICBM152-2009c MNI space via combinations of linear and diffeomorphic transformations. The brain was seamlessly reconstructed from the fields of view with q-space reinterpolation to merge all the dMRI in the same q-space sampling. (C) Local diffusion models, including DTI and NODDI or MWF mapping, were subsequently applied to unveil intricate details.

### Touring the human brain at the mesoscopic scale

Ultra-high resolution, coupled with the various contrasts available in anatomical, diffusion, and quantitative MRI, enabled the identification of many anatomical structures inaccessible at lower fields (Fig. 4). For instance, the basal ganglia could be segmented with great precision. Their substructures could be identified by combining contrast mechanisms from aMRI, qMRI, and dMRI, thereby enhancing the accurate delineation of substructure boundaries. The high iron content of specific structures has an impact on T2* contrast, allowing for the delineation of various brainstem nuclei such as the substantia nigra, the subthalamic nucleus, or the red nuclei, but also the amygdala, whose substructures were shown to exhibit some T2* hyperintensities related to ageing (*39*). Quantitative R2* maps also revealed hyperintensities in superficial white matter (SWM) bundles, where myelinated axons are known to embed higher iron content in subcortical regions (*14*).

**Fig. 4:**
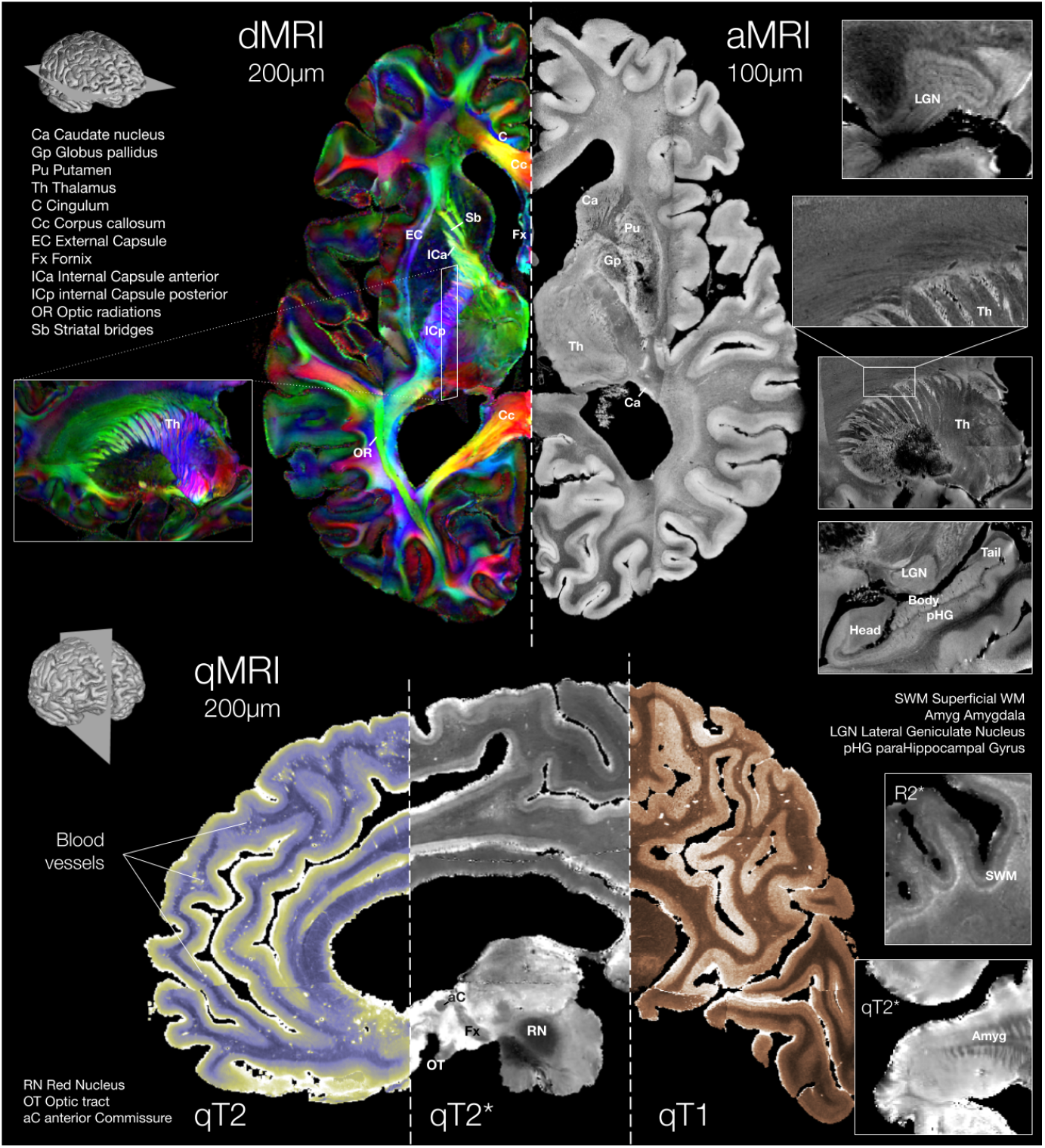
dMRI, aMRI, and qMRI maps of the Chenonceau brain. Top: main direction colour-encoded map stemming from the DTI model, and highly-resolved anatomical map at 100 μm, along with focus on the thalamus (Th) and its connections, the layers of the lateral geniculate nucleus (LGN) revealed at such high resolution, and the hippocampus fine anatomy shown in sagittal view. Bottom: qT2, qT2*, and qT1 quantitative maps at 200 μm. The high mesoscopic resolutions allowed for the identification of subcortical nuclei, including the caudate nucleus (Ca), the globus pallidus (Gp), the putamen (Pu), the thalamus (Th), the red nucleus (RN), as well as, white matter fibre bundles, including the cingulum (C), the corpus callosum (Cc), the external capsule (EC), the fornix (Fx), the internal capsule anterior (ICa) and posterior (ICp), the optic radiations (OR), the striatal bridges (Sb), the optic tract (OT) and the anterior commissure (aC). qT1 and qT2 contrast differences observed between blood vessels and brain parenchyma contribute to highlighting the vascular tree. Fine mesoscopic details of structures known to embed a high iron content, such as the amygdala (Amyg), shown in axial view, or the SWM, are revealed through the observation of signal hypointensities in R2* maps.

Although blood was flushed from the vascular compartment by the formalin intra-arterial infusion, the mesoscopic vascular tree remains particularly visible on T1- and T2-weighted volumes due to the substitution of blood with buffered formalin solution in the deepest vessels.

Anatomical imaging at 150 μm has a higher gray-white matter contrast and a lower sensitivity to residual magnetic susceptibility effects. However, the 100 μm anatomical imaging sequence offers higher resolution and provides further detail on the smallest anatomical structures of the midbrain and cerebral cortex. It revealed structures never observed before from MRI, such as the lateral geniculate nuclei layers, alternatively connected to the ganglion cells from both retinas involved in the foveal, high-resolution, lower sensitivity, and color-sensitive perception (parvocellular/midget cells), and in the peripheral, low resolution and higher sensitivity, black and white perception (magnocellular parasol cells) (Fig. 4).

The hippocampi, playing a central role in episodic memory, could also be identified with unequaled precision, segmentable in subsequent steps, and discriminated by their subfields using T2 and T2* contrasts and neurite density. The Chenonceau dataset offers the opportunity to study its two hippocampi, their subfields, microstructure, and connections, which allows direct comparison between the two homologous allocortical structures, a possibility rarely offered in other studies (*34–35*).

### Exploring the nested connectome of the Chenonceau brain

A tractogram comprising approximately 25 million streamlines was computed using a dedicated high-performance computing global tractography reconstruction framework relying on a spin-glass model to robustly deal with complex local fiber configurations and to address the need for a high-resolution dMRI dataset while preserving computation time (*40, 41*). Using a well-established segmentation approach relying on a novel deep white matter (WM) atlas (*42*), fifty deep WM fiber bundles were recovered with high-precision trajectories (Fig. 5A), including complex connectivity patterns such as crossing bundles between thalamic and lenticular radiations or 90° sharp turns of frontal thalamic radiations when exiting the thalamus. The large number of shorter connections necessitated additional dimension reduction steps to facilitate efficient classification and identification of connections. To this end, a tractogram analysis using unsupervised clustering methods (*43*) identified more than 2,000 short association WM fiber bundles per hemisphere. The reconstructed short bundles could be further split into two categories based on length, yielding subcortical (Fig. 5B) and intracortical (Fig. 5C) connections. To further illustrate subcortical connectivity, bundles surrounding the central sulcus were extracted with high precision, highlighting the many symmetrical connections between pre- and post-central gyri. Intracortical connections gathered an outstanding set of very short U-shaped bundles densely distributed over the cortex.

**Fig. 5:**
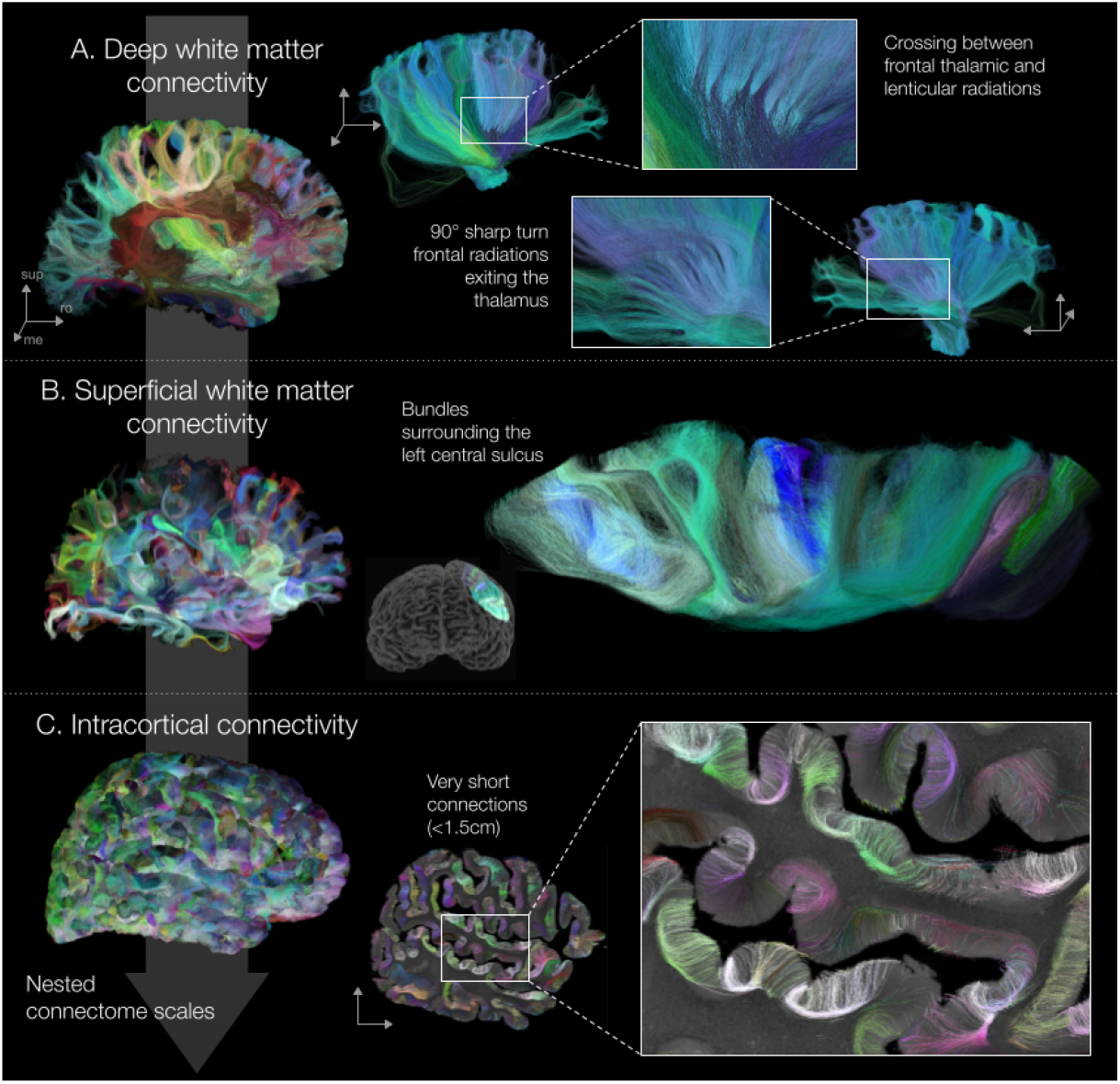
Nested connectome of the Chenonceau brain, from deep WM connections (A), revealing fine details, to subcortical connectivity (B) and intracortical connectivity (C). (A) Identification of the thalamic and lenticular radiations of the right hemisphere, along with the corticospinal tract, on the lateral-to-medial view highlights their crossing. The medial-to-lateral 3D rendering highlights the 90° sharp turns of the fiber trajectories exiting the thalamus. (B) 3D rendering of the SWM bundles surrounding the central sulcus, whose trajectories were reconstructed with a high level of accuracy. (C) Observation of the shortest connections (fiber length < 1.5 cm) reveals the intracortical connectivity of the Chenonceau brain, reconstructed from mesoscopic-resolution diffusion MRI data. All fibers are color-encoded according to their tortuosity vector, exacerbating the directionality of SWM and intracortical connection curvature (see Methods for further explanation).

### Mapping the cortical laminar structure

The multimodal approach enabled in-depth phenotyping of the Chenonceau cortex microarchitecture using relevant qMRI and dMRI-based markers of cyto- and myeloarchitecture (*20, 12*). The high resolution of qMRI and dMRI data allows for in-depth exploration of the cortical ribbon and detailed characterization of quantitative parameters derived from these modalities within the cortex (T1, T2, and T2* relaxation times, as well as myelin water fraction for qMRI, and neurite density and orientation dispersion of dendrites for dMRI), whose visual inspection highlights the local laminar structure of the cerebral cortex (Fig. 2). A compressed voxel-wise microstructural signature representation was learned from a variational autoencoder (VAE), trained on a set of MRI microstructural features within the cortical ribbon of the Chenonceau brain. While 2 latent dimensions showed high tangential variability, 2 others showed mostly radial cortical variability (Fig. 6). These latter were used in a classification task proposed as an extension of the approach in (*44*) to ultimately expose the layered organization of the Chenonceau cortex (Fig. 6).

**Fig. 6:**
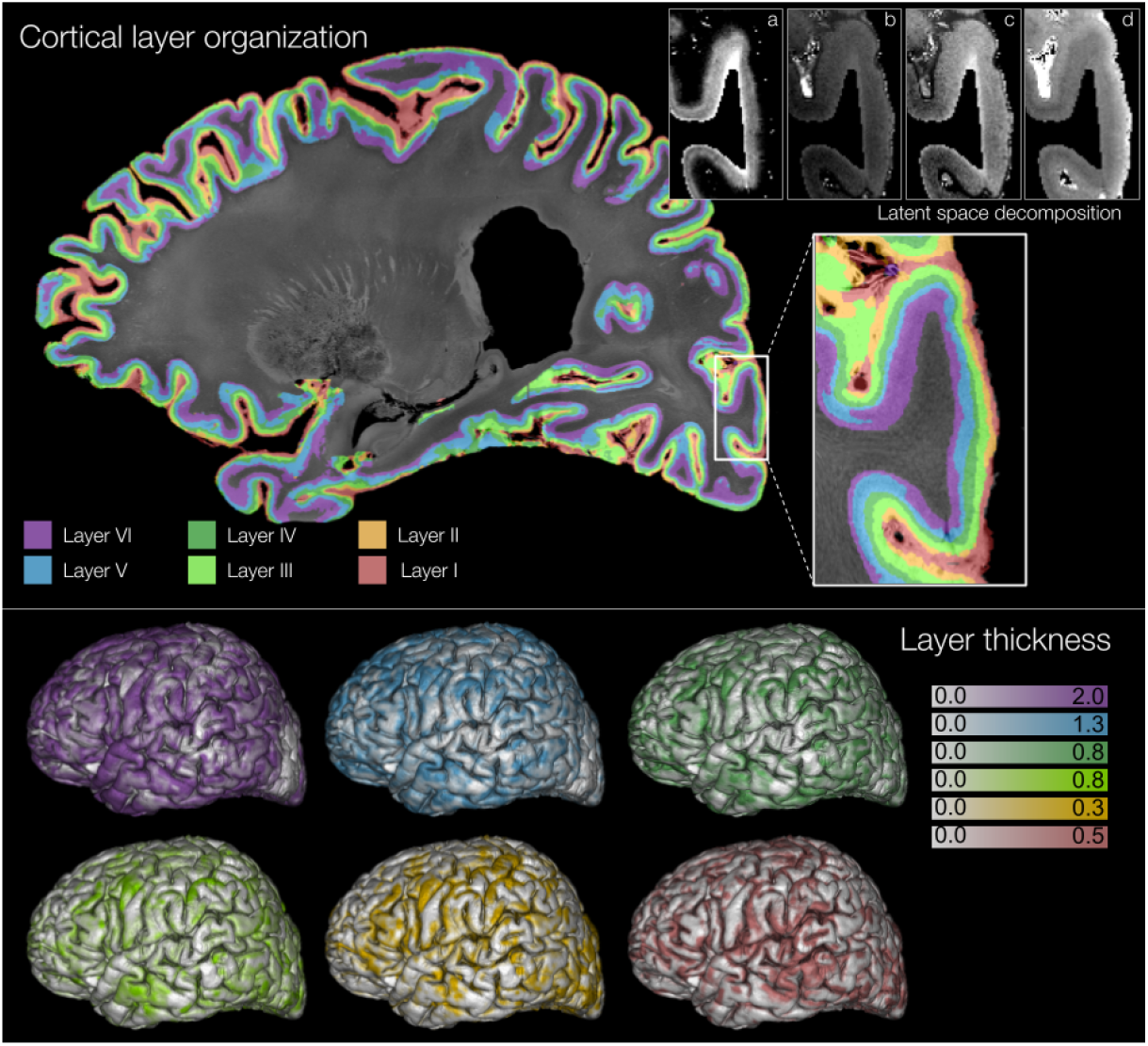
Relaxometric and diffusion quantitative MRI maps revealing the cortex laminar structure. Upper right corner: the four latent dimensions deduced after VAE training in a region corresponding to the visual cortex. Latent dimensions (a) and (d) were used to fit a Gaussian mixture model (GMM) in order to recover the layered structure (top) at a super-resolution of 100 μm. Bottom: the distributions of layer thickness at the brain surface show high tangential variability in the identified layers.

## Discussion

Reconstructing an entire volume from *post-mortem* samples after an irreversible cutting step remains an unsolved challenge; the lack of overlap between blocks is the main obstacle to accurately reconstructing the whole brain from individual blocks. A few recent studies have addressed this issue in the literature (*45, 46*), and these mainly focus on histology, which requires sub-sampling. For the Chenonceau brain, the method proposed in (*47*) may not yield satisfactory reconstructions, as it relies on linear transformations. In this work, we developed a new methodology that successfully integrates nonlinear transformations into the reconstruction process to account for deformation sources (including cutting, storage, or rehydration of the samples).

While tractography inherently suffers from biases at the millimeter scale (*48*), diffusion acquisitions with high spatio-angular resolutions are expected to mitigate many of these limitations. The high diffusion sensitivities (up to 8000 s.mm^-2^) yielding high angular resolutions, combined with the high spatial resolution of the Chenonceau dMRI dataset, considerably enhanced the estimation of the direction of axon populations during fiber tracking. The reduction in partial-volume effects, enabled by the 200 μm resolution, revealed the expected sharp turns at the gray/white matter interface and significantly contributed to reducing the gyral bias (*49*). Furthermore, the high-resolution aMRI, qMRI, and dMRI maps provided key information, further enhancing the robustness of global tractography methods for accurately reconstructing the trajectories of fibers entering the cortical mantle and their terminations.

The complete mesoscopic mapping of the superficial connectivity enabled the virtual dissection of macroscopic connectivity structures that are only partially observed at millimetric resolutions and remain controversial. For instance, a focus on the central sulcus has revealed the many symmetrical connections between the primary and the secondary motor cortices, with fibers converging centrally, corroborating recent inside-out observations of short association fibers in Klingler’s dissection (*50*). The identification of omnipresent intracortical connections suggests highly localized and directionally coherent communication patterns between neighboring cortical areas. While these connections remained intractable in standard dMRI datasets, similar ones have already been observed, yet not systematically reported, in anatomic tracing studies (*51–53*).

The qMRI protocol aimed to explore quantitative markers of myeloarchitecture, such as the myelin water fraction, and of iron load within structures. To pave the way for more complex models, a deliberate decision was made to collect a large number of T1-, T2-, and T2*-weighted samples for this study. This allowed the robust inference of myelin markers across the entire Chenonceau brain from a multi-compartment model. As shown in Fig. 2, the MWF map reveals periventricular WM areas with severe myelin loss. These areas consistently correlate with the hyperintensities observed on the T2-weighted map, commonly associated with demyelination or axonal degeneration (*54*), which may originate from vascular dysfunctions common in such elderly subjects.

Previous studies focusing on the development of multi-compartment T1-weighted signal models combined with clustering techniques have already demonstrated their potential for identifying cortical layers (*55, 56*). Diffusion MRI extracted features (*57, 58*) are believed to be less robust for performing this task (*59*). However, in the Chenonceau dataset, the approach was extended to incorporate multimodal descriptors derived from qMRI and dMRI, covering both myeloarchitectural and cytoarchitectural information. Further investigations could be conducted to assess the coherence of the laminar structure established in this study with that reported in histology (*1–3, 60*). While the classification method was first applied to the radial direction to segment the laminar structure, this highlights the need to extend the classification approach over the cortical surface to capture the cyto-/myelo-architectonic tangential variability. It would ultimately enable the production of novel architectonic areas similar to those observed in histology (*61*), paving the way for a comprehensive multimodal cortical parcellation of the Chenonceau brain, concomitant with its fine structural connectivity.

The multimodal ultra-high-field MRI acquisition campaign offers the opportunity for in-depth phenotyping of an entire *post-mortem* human brain at the mesoscale. This effort yields a novel foundational dataset that: 1) establishes a new whole high-resolution reference cerebrum, 2) bridges the scale gap between *in vivo* imaging and *post-mortem* microscopic datasets, and 3) promotes the joint analyses at the mesoscale of fine anatomical structures, their structural connectivity, vascularization, and microarchitecture to establish new models of the human brain. The reconstructed Chenonceau dataset is available on the open-data EBRAINS portal (https://www.ebrains.eu/), both in its native space and in the stereotaxic MNI ICBM 2009c nonlinear asymmetric space.

## Methods

### Tissue preparation

Thirty-two hours after death, 4% buffered formalin was injected into both internal carotid arteries for optimal *in calvaria* fixation (*62*). It was extracted 24 hours later and remained immersed in the same solution for an additional five months, hung by the basilar artery to prevent deformation induced by contact with the container.

After a 3-month rehydration step in 0.1M phosphate-buffered saline (PBS) solution, the cerebrum was separated from the rhombencephalon at the base of the midbrain and sagittally split on the midline. To prevent deformation, the ventricles were filled with an MRI-invisible gel selected for its magnetic properties, stability over time, and ease of processing at room temperature (Biz’gel sealing gel, BizLine, France). Each hemisphere was further cut into two 4 cm-thick slices in the medio-lateral direction and subsequently divided into three or four blocks in the dorso-ventral direction. This process resulted in 13 parallelepipedal blocks with a maximum span of 20 cm to cover the entire cerebrum length in the rostro-caudal direction. These blocks were placed in gel-filled 3D-printed plastic containers to immobilize them, ensuring reproducible positioning in the MRI scanners between imaging sessions (see supplementary Fig. 8). All blocks were preserved at a constant temperature of 5°C.

A specific imaging protocol was designed on the 3 Tesla Prisma MRI scanner (Siemens, Erlangen) for blockface and blocks imaging, including two T2-weighted imaging SPACE sequences: before cutting to obtain a blockface image, echo time T_E_ = 108 ms, repetition time T_R_ = 700 ms, isotropic resolution of 500 μm, one average, readout bandwidth RBW = 200 Hz/pixel; and after cutting to provide intermediate reference images for reconstruction, T_E_/T_R_ = 103/700 ms, isotropic resolution of 400 μm, four averages, RBW = 200 Hz/pixel.

### Ultra-high field MRI protocols

At 11.7T, the sensitivity profile of the volume coil antenna constrained the acceptable field-of-view length along the magnet axis to the 60 mm inner diameter of the coil. Further investigation of the image quality led to selecting a FOV of 42×42×56 mm^3^ for all sequences, with two additional 7-mm extensions along the magnet axis to facilitate consecutive FOV matching with overlapping regions.

#### aMRI protocol at 11.7 Tesla

The aMRI protocol included two anatomical scans using 2D and 3D spin-echo pulse sequences, chosen for their innocuousness to magnetic susceptibility effects on a relatively large field-of-view, especially at 11.7T. The parameters of the 2D spin echo imaging sequence of isotropic spatial resolution of 150 μm are the following: T_E_/T_R_ = 16/6647 ms, field of view FOV = 41.25×41.25 mm^2^, matrix size 275×256, slice thickness = 150 μm, number of interleaved slices = 374, nine averages, flip angle = 90°, RBW = 66 kHz, total acquisition time 3h26min. The parameters of the 3D spin echo imaging sequence of isotropic spatial resolution of 100 μm are the following: echo time T_E_/T_R_ = 20/500 ms, field of view FOV = 40×40×56 mm^3^, matrix size 400×400×560, number of averages N = 1, flip angle = 90°, RBW = 50 kHz, total acquisition time = 22h13min.

#### dMRI protocol at 11.7 Tesla

The diffusion-weighted MRI dataset was acquired using a multiple shell sampling of the q-space over three spheres at b = 1500/4500/8000 s.mm^-2^. The number of diffusion directions uniformly distributed on the surface of each sphere was set to 25, 60, and 90 directions, respectively. A 3D segmented EPI diffusion-weighted Pulsed Gradient Spin Echo (PGSE) sequence was chosen to keep the total scan duration acceptable for a single field of view, with the following parameters: isotropic spatial resolution of 200 μm, T_E_/T_R_ = 24.3/250 ms, diffusion pulse width = 5 ms, diffusion pulse separation time Δ = 12.3 ms, one average, FOV = 42.4×40.8×56.0 mm^3^, matrix size 212×204×280, flip angle 90°, RBW = 300 kHz, 30 segments, number of reference volumes at b = 0 s.mm^-2^ set to N_b=0_ = 17, total acquisition time = 82h. The non-standard acquisition size on this MRI, typically used for rodent imaging, necessitated splitting the dMRI acquisition into 17 scans so that the raw data from each could fit into RAM before reconstruction. 10-minute pauses were added after each scan to maintain the sample temperature at equilibrium throughout the 82h of acquisition, which is fundamental for exploring the diffusion process, known to be temperature-dependent.

#### qMRI protocol at 7 Tesla

T1-weighted Variable Flip Angle (VFA) Spoiled Gradient Echo (SPGR) volumes were collected using 3D VFA FLASH sequences (*63*): T_E_/T_R_ = 4.99/15 ms, 65 flip angles, sampled every 0.5° between 0° and 30°, then every 1.5° between 30° and 45°, isotropic spatial resolution of 200 μm, matrix size = 280×212×204, 1 average, RBW = 50 kHz, scan time = 11h43min. T2-weighted Multi Spin Multi Echo (MSME) volumes were collected using a 3D MSME sequence: T_E_/T_R_ = 5.56-166.8/1000 ms, 30 echoes, echo spacing = 5.56 ms, isotropic spatial resolution of 200 μm, matrix size = 280×212×204, three averages, RBW = 100 kHz, scan time = 36h02min. T2*-weighted 3D FLASH volumes were collected with 3D variable echo time FLASH SPGR sequences: T_E_/T_R_ = 5-100/110.048 ms, 10 echoes sampled every 5 ms between 5 and 20 ms and every 10 ms between 20 and 100 ms, flip angle = 50°, isotropic spatial resolution of 200 μm, matrix size = 280×212×204, one average, RBW = 50 kHz, scan time = 13h10min. Additionally, a dual flip-angle 3D EPI B1+ mapping sequence was acquired in order to estimate the B1 inhomogeneity field (*64, 65*): T_E_/T_R_ = 13.405/1500 ms, flip angles = 30/60°, 1 average, 35 segments, signal type = spin echo, isotropic spatial resolution of 200 μm, matrix size = 280×212×204, RBW = 300 kHz, scan time = 5h56min. Fat saturation pulses were applied to all sequences to effectively suppress the gel signal without affecting the tissue signal. Dummy scans were added to the protocol, as in the 11.7T acquisition protocol, to reach magnetization equilibrium and produce a homogeneous signal across the scanned FOV. Pauses of ten to twenty minutes were inserted between each series of MRI sequences to preclude the brain block from heating during acquisition.

#### Acquisition campaigns

The blocks were stored at 5°C in phosphate-buffered saline. One day before the acquisition, PBS was replaced with Fomblin Y LVAC 06/6 (perfluoropolyether, Solvay Specialty Polymers USA, LLC, West Deptford, NJ), which is invisible to MRI and reduces magnetic susceptibility artifacts. During this procedure, air bubbles that may have been trapped in the sulci were manually eliminated. The sample was left at room temperature for at least 12 hours (*66*) before acquisition. Since each block had to be acquired multiple times, an accurate and reproducible positioning in the scanner was essential. To achieve this, we used the AutoPac motorized table of the Bruker 11.7 T scanner, which offers sub-millimeter accuracy. The sample then underwent its target imaging protocol, lasting 107h to scan a 42×42×56 mm^3^ field of view within the block on the 11.7 Tesla Bruker MRI system and 66h to scan the same field of view on the 7 Tesla Bruker MRI system. After each acquisition, Fomblin was replaced with PBS before being stored at 5°C until the next acquisition session. Collecting the mesoscale aMRI, qMRI, and dMRI datasets across 45 FOVs covering 13 tissue blocks took almost 2 years on the NeuroSpin 7T and 11.7T preclinical MRIs (see supplementary Fig. 9).

### Reconstruction of the whole brain

The strategy used to reconstruct the Chenonceau brain consisted of 1) aligning the mesoscopic FOVs to their corresponding block and 2) reassembling the entire brain from the blocks using the intermediate scans made at 3T. Developed in a parallelized context, the registration pipeline involved the computation of 464 3D diffeomorphic transformations using ANTs software (*67, 68*) to account for the multiple sources of deformation between acquisitions at 3, 7, and 11.7T.

#### Intermediate blocks reconstruction

First, all multimodal FOV volumes were non-linearly aligned to predefined reference spaces in order to account for non-aligned FOV centers across acquisition sessions, using high-SNR reference volumes with contrasts similar to those of T2-weighted images at 3T. These reference FOVs were then aligned to their corresponding 3T blocks using linear and diffeomorphic transformations, initialized with preliminary translations along the block axis. Additional full-scale diffeomorphic registrations were performed on overlapping areas between consecutive FOVs to further match the intricate details of the mesoscopic data, similarly to (*69*). The high-resolution blocks were then recomposed based on a criterion that combined voxels when FOVs overlapped to preserve signal integrity within voxels. For dMRI, this criterion was based on the maximum intensity signal among the different overlapping areas of the volumes at b = 0 s.mm^-2^, assuming that lower intensity values correspond to regions of weak coil sensitivity. Finally, in the same block, all modalities, with their respective reference contrasts, were co-registered to the 200 μm T2-weighted, TE = 22.24 ms block, chosen as the reference, as its histogram shape matched those of the acquisitions performed at 3T.

#### Whole brain reconstruction

The inverse problem of reconstructing the 3D brain from cut blocks was addressed by 1) defining a target region within the blockface for each block, and 2) computing independent linear and diffeomorphic co-registration of each block to its target. To define these targets, an iterative mask delineation strategy was implemented to subdivide the blockface into non-overlapping, gap-free block regions consistent with the blockface geometry. The developed approach was motivated by the empirical observation that repeating the diffeomorphic registration process with increasingly fine segmented targets from previous iterations improved the overall registration accuracy. At initialization, independent unconstrained block-to-blockface registrations were computed, preliminary initialized by manual linear positioning, yielding initial block masks. Iteratively, these masks were combined using geodesic distance criteria to split overlaps and fill gaps between neighboring blocks. It was followed by diffeomorphic co-registrations using the updated targets and relaxed constraints on blocks’ spatial extent to allow exploration, yielding refined masks for each block extent. Iterations were first performed using the 400 μm blocks acquired at 3T and then, using the 200 μm T2-weighted (T_E_ = 22.24 ms) blocks. The obtained subdivision was then used as domain constraints in the final registration process to ultimately match the 13 blocks to the blockface. The blockface volume was then brought back into the MNI ICBM152 Non-Linear Asymmetric 2009c anatomical template space following a classical diffeomorphic normalization task.

#### Recomposition of the whole brain

Supplementary Fig. 10 summarizes the transformation paths bringing the high-resolution FOVs into the MNI space. Each path corresponds to the composition of 4 to 6 consecutive transformations. The whole brain was reconstructed at the mesoscopic level, composing the set of transformations in a single interpolation step (spline 3^rd^ order). dMRI data were also reinterpolated in q-space to account for diffeomorphism-induced local rotations. A target set of directions was specifically set for the three shells, keeping the same number of directions. A re-interpolation step of the diffusion-weighted signal in the q-space over each of the three spheres was done: diffusion signal was transformed into the spherical harmonics basis (*17*), from which reoriented dMRI data were recovered by the inverse transform towards the target q-space while taking into account the full path of local rotations. Local diffeomorphism-induced rotations were computed as proposed in (*70*).

#### Signal intensity homogenization

Due to different scaling factors at the digitization, reconstruction, and preprocessing stages of the aMRI, qMRI, and dMRI datasets, the reconstruction pipeline had to consider the non-quantitative nature of the MRI signal. To this aim, FOVs were scaled at recomposition based on histogram matching. Boundary effects, due to the bias field at the FOV’s extremities, were corrected by integrating a continuity constraint when accounting for the bias field. The continuity constraint was implemented via a normalization factor α of the low-frequency signal within each individual FOV mask and the low-frequency signal over the whole-brain mask. Low-frequency components were obtained by convolving the image with a 3D Gaussian kernel *G* (σ = 1.5 mm):

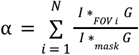

where *I* is the image, denotes ∗ _*FOV i*_ the convolution operation restricted to the mask of FOV *i*, and ∗_*mask*_ denotes the convolution performed over the whole-brain mask.

### Inference of structural connectivity

An accurate mask of the cerebrum was first computed from T1 and T2 qMRI maps using edge-detection methods and manual delineation. The b = 8000 s.mm^-2^ diffusion dataset was used to compute an orientation distribution function (ODF) map using the analytical Q-ball model (*17*). The maximum harmonic order was set to 8, and the Laplace-Beltrami regularization factor to 0.006. Global tractography methods (*40*) retain a probabilistic nature and address the shortcomings of streamline tractography methods by competing the creation of fibers in order to achieve an optimal solution to the ill-posed problem of inferring fiber trajectories from an ODF map. A global approach based on a spin glass model was therefore chosen for its robustness to the creation of false positives. The ExaTract global tractography tool, relying on a distributed implementation and consequently compatible with high performance computing facilities (*41*) was used to infer the structural connectivity of the Chenonceau brain from the former ODF map within the precomputed tractography domain with the following parameters: whole brain subdivided into 800 parcels, four initial spin-glasses per voxel of length 160 μm, relative probabilities of connection/motion/creation/deletion ratio = 1/4/0.8/0.05, 3 simulated annealing cycles of initial/final temperatures 0.1/0.003K, 0.05/0.003K, and 0.025/0.003K respectively, a connection likelihood of 0.5 and energy weighting α_connexion_/□_ODF_ = 4.0/5.0. The tractography was run on 100 nodes of the Irène-Rome supercomputer (corresponding to 12,800 CPU cores; Très Grand Centre de Calcul, CEA DAM-Ile-de-France, Bruyères-le-Châtel).

#### Bundle segmentation

Deep WM bundles were segmented using a fiber labelling approach (*43*) based on a deep WM atlas recently published and established at the millimeter scale from the entire HCP cohort (*42*). For the segmentation of shorter WM bundles, we used an unsupervised approach and applied a hierarchical fiber clustering algorithm (43) to reconstruct fiber clusters without prior knowledge from fibers shorter than 85 mm. This parcellation-based clustering method was used with the following parameters: a parcellation resolution of 2 mm, a fiber range of 10 mm, regularly spaced every 5 mm, a connectivity matrix threshold of 0.0005, and a minimum percentage of fiber length intersecting a parcel cluster fixed at 20%. The obtained clusters were further aggregated to establish WM bundles using an HDBSCAN clustering algorithm over the normalized pairwise distances between cluster centroids (*43*). Parameters were set to: fiber normalization factor of 14 mm, HDBSCAN neighbor count of 40, and minimum cluster size of 1, percentage of cluster occurrence in the population of 0% as a single subject.

#### Fiber color-encoding scheme

For visualization purposes, the fibers shown in Fig. 5 are represented with an HSV color-encoding based on their tortuosity vector, as proposed in (*71*). Unlike conventional color-encoding based on fiber local directionality, this encoding accentuates the shape and orientation of the fibers. Initially intended for long WM tracts, this color-encoding has proven particularly well-suited to U-shaped bundles by accentuating the direction of their curvature (see supplementary Fig. 11).

### Computing proxies to myelo- and cyto-architecture

#### Quantitative relaxometry times

Maps of quantitative T2 and T2* relaxation times were estimated voxel-wise from a log-linear regression based on a Levenberg-Marquardt optimizer. The map of quantitative T1 relaxation time was estimated using a non-linear programming (NLP) optimizer accounting for the Rician noise corrupting the data.

#### Myelin water fraction mapping

Largely adapted from models in (*12*), assuming a “fast exchange” model of water molecule (*72*), the T1-weighted signal resulting from the contribution of all these compartments can be modeled with an apparent T1_ap_ relaxation time 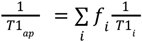, where *f*_*i*_ is the volumic fraction of compartment *i* such that 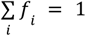. The corresponding T1-weighted signal contrast measured at each voxel using a VFA FLASH pulse sequence can be modeled by the equation:

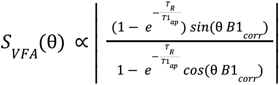

where θ is the flip angle, and *B1*_*corris*_ the B1 inhomogeneity correction map. The T2-weighted transverse relaxation is commonly considered to follow a “slow exchange model” (*72*), yielding a simple linear composition of the contributions of the two or three chosen compartments. The corresponding T2-weighted MSME signal can then be modeled by the following equation:

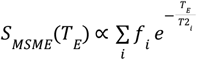

To identify the compartments and robustly infer their magnetic properties, an initial 3-compartment model analysis was conducted on a sub-part of the brain (FOV LC1) without any constraint except that the sum of their corresponding volume fractions must be equal to 1. The model was fitted using a non-linear programming (NLP) optimizer, accounting for the Rician noise. The resulting histograms of the T1 and T2 relaxometric times revealed the existence of three distinct compartments. In comparison with (*12*) and in the absence of CSF, the three compartments were attributed to myelin-related protons, white matter protons, and gray matter protons, with respective relaxometric times: T2_my_,= 4.5 ± 3 ms, T1_my_ = 235 ± 70 ms, T2_WM_ = 30 ± 8 ms T1_WM_ = 230 ± 70 ms, T2_GM_ = 56 ± 10 ms, T1_GM_ = 285 ± 70 ms. To robustly compute the myelin compartment fraction throughout the brain, the 3-compartment analysis was then performed voxel-wise, assuming fixed relaxation times chosen to correspond to the peak values inferred from the precomputed qT1 and qT2 histograms, as determined in the previous step.

#### Diffusion tensor imaging

A DTI map was computed from the diffusion-weighted MRI dataset at b = 4500 s.mm^-2^ to provide quantitative scalar maps of fractional anisotropy, apparent diffusion coefficient, parallel and transverse diffusivities, as well as a color-encoded direction map.

#### NODDI model

The microarchitecture of neurites populating both the white matter (with fibers) and cortex (with dendrites and few fibers) was evaluated in the Chenonceau brain using the microstructural NODDI model (*20*) providing access to the volume fraction of neurites within each voxel of the brain, as well as their orientation dispersion, assuming a Watson distribution of the neurite and a hindered Gaussian distribution to represent the extra-axonal space. Using the signal stemming from the three shells, fixed parallel diffusivity set at 3.0 × 10^-10^ m^2^/s and fixed isotropic diffusivity set at 2.0 × 10^-9^ m^2^/s (estimated from the diffusivity obtained from the DTI model), the model was fitted voxel-by-voxel using an NLP optimizer (with a maximum iteration count of 1000 and a maximum error of 0.01).

### Cortical layer segmentation

The cortex was segmented from a preliminary Gaussian mixture model run with 2 clusters on selected maps, which showed the best contrasts between gray and white matter at a 200 μm resolution. The neurite volume fraction map from the NODDI model and the proton density map derived from T2-relaxometry mapping were particularly well-suited for this task. The obtained density probability map was thresholded to delineate the cortex. Deep gray matter nuclei were excluded based on ROIs drawn manually at lower resolutions. The resulting cortex mask finally contained approximately 71 million voxels.

Voxel-wise vectors of microstructural signatures were built from the 200-μm-resolution features, including qT1, qT2, qT2*, neurite orientation dispersion, and neurite volume fraction. Outliers were first removed using unsupervised outlier detection based on the Local Outlier Factor (LOF). To reduce the input feature vector for the GMM classifier and select the adequate features, a non-linear data compression was performed based on a VAE (*73*). This results in a latent feature vector of dimension 4, sufficiently dense for the classifier model to robustly estimate the Gaussian distributions. From these four latent maps, only two were selected based on their visual contrast in accordance with the radial organization of the cortical layers.

The Gaussian mixture model was fitted with data stemming from the two selected latent dimensions. To obtain an adequate number of target clusters, several configurations were tested and compared based on the Akaike Information Criterion (AIC) and the Bayesian Information Criterion (BIC). The configuration yielding 22 clusters was selected, achieving the lowest AIC and BIC values. Voxels were assigned to clusters based on their maximum posterior probability. Among the 22 clusters, 14 clusters were not representative of the voxel population (less than 0.5% of the total number of voxels).

They mostly represent incoherent NMR signatures of voxels found with partial volume outside the brain that were included in the cortex mask previously obtained. These clusters were merged with the cluster representing the outer layer. Without any additional prior knowledge, the 8 clusters successfully reconstruct some distinct microstructural layers within the cortex. The inner one was thought to represent WM or partial volume with WM. The two outermost clusters were merged into a single cluster, as their spatial extents alternated.

While VAE training and GMM fitting were performed on the MRI data at their native resolution of 200 μm, the final layered map was inferred at the super-resolution of 100 μm based on reinterpolated qMRI and dMRI maps further processed with the trained VAE in order to provide smoother geometric boundaries between layers. The visual consistency of these boundaries was further improved by a Potts regularization model. The label *u*_*i*_ at voxel i is established by a trade-off between its original value and the labels of its neighbor voxels, such as:

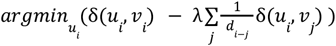

where *d*_*i*−*j*_ is the Euclidean distance between the centers of voxel *i* and of voxel *j, v*_*i*_ is the label of voxel *i*, and δ is the Dirac function. 1.2×10^10^ regularization iterations were performed on the labeled layer map at 100 μm resolution, using a neighborhood containing the 26 nearest voxels and a regularization factor of 2.0.

## Acknowledgments

The authors warmly thank Dr. Fawzi Boumezbeur, Dr. Françoise Geffroy, and Jeremy Bernard for their technical support for the various tasks involving the 11.7 Tesla MRI platform, the histology lab, and the mechanical lab, respectively. The authors sincerely thank those who donated their bodies to science so that anatomical research could be performed. Results from such research can potentially increase humankind’s overall knowledge, which can then improve patient care. Therefore, these donors and their families deserve our highest gratitude. The authors would also like to thank Dr. France Boillod-Cerneux and Dr. Christophe Calvin for their support within the framework of the French-German AIDAS institute, as well as the GENCI French national HPC infrastructure.

## Funding

French National Agency, FibrAtlas-II-III, ANR14-CE17-0015-01 (CD)

European Union’s Horizon 2020 Framework Programme for Research and Innovation, Specific Grant Agreement No. 945539 (Human Brain Project SGA2 and SGA3) (CP)

Joint German-French Institute AIDAS Cycle 1 and Cycle 2

Computational resource GENCI allocation A0120310851 and GENCI special allocation SS010315366 on the Irène-Rome partition of the Joliot-Curie supercomputer (TGCC, Bruyères-le-Châtel, France)

MRI acquisition fully covered by the NeuroSpin imaging platform

## Contributions

Conceptualization: CP, CD ; Investigation: JB, RYH, AP, CP, CD, ILM ; Methodology: SL, AP, CP ; Software: SL, CP, FP, AP ; Formal analysis: SL, BH ; Visualization: SL ; Funding acquisition: CP, CD; Supervision: CP, CD, IU ; Validation: AP, SL ; Data curation: SL, IU ; Writing – original draft: SL, CP ; Writing – review & editing: SL, CP, CD, ILM, IU

## Ethical considerations

The brain was obtained from the body donation program of Université de Tours. Before death, participants gave their written consent for using their entire body – including the brain – for any educational or research purpose in which the anatomy laboratory is involved. The authorization documents (in the form of handwritten testaments) are kept in the files of the Body Donation Program.

## Competing interests

The authors have no competing interests to declare.

## Data availability

The reconstructed dataset is available on the open-data EBRAINS portal (https://www.ebrains.eu/), in the native space of the Chenonceau brain and the stereotaxic MNI ICBM 2009c nonlinear asymmetric space. Scalar maps, deep connections, short connections, and cortical layer mapping are available in the following datasets: https://doi.org/10.25493/SJP6-YW0, https://search.kg.ebrains.eu/instances/494818fa-9126-4432-a167-52ff02534a8b (currently being curated), https://search.kg.ebrains.eu/instances/657fc3ae-9e65-44eb-99ca-7e92b4ab4b26/ (currently being curated), and https://search.kg.ebrains.eu/instances/cbf3220a-524a-4473-96b8-fc9dcb0f55fe (currently being curated).

## Code availability

FOV preprocessing (including qMRI and dMRI) and recomposition processes were performed with the Ginkgo toolbox, which is freely available at https://framagit.org/cpoupon/gkg. ANTs was used to perform registration processes and N4 bias field correction. The VAE and the Gaussian Mixture Model were implemented using PyTorch and the scikit-learn library, respectively.

## Notes

https://doi.org/10.25493/SJP6-YW0

https://search.kg.ebrains.eu/instances/494818fa-9126-4432-a167-52ff02534a8b

https://search.kg.ebrains.eu/instances/657fc3ae-9e65-44eb-99ca-7e92b4ab4b26/

https://search.kg.ebrains.eu/instances/cbf3220a-524a-4473-96b8-fc9dcb0f55fe

### Competing Interest Statement

The authors have declared no competing interest.

