## Supplementary material for "Chenonceau: mesoscopic MRI deep phenotyping of a post-mortem human brain at 7 and 11.7 Tesla"

Supplementary text 1

**Quality check and SNR measurements**

To check the quality of the data and to assess the stability of the imaging protocol and tissue over time, a systematic evaluation of the signal-to-noise ratio (SNR) of the various anatomical and diffusion datasets collected at 11.7 Tesla was performed. Regions of interest corresponding to white matter were semi-automatically defined within the tissue across the different fields of view. Each region of interest was carefully chosen to correspond to a homogeneous tissue region. The regions were also selected near the center of the field of view to reduce the impact of the antenna intensity profile.

The SNR was evaluated in each region by measuring the ratio between the average signal obtained within the region and the standard deviation of the noise measured in a mask corresponding to the image background. Fig. 7A illustrates the SNR measurements obtained for all anatomical scans, all fields of view, and all blocks. The average SNR was estimated at $SNR_{Anatomy(100\mu m)} = 36.7 \pm 11.8$ for the anatomy scans at 100 µm resolution, and at $SNR_{Anatomy(150\mu m)} = 17.5 \pm 4.2$ for the anatomical scans at 150 µm resolution. Fig. 7A also illustrates the SNR values obtained for diffusion-weighted MRI scans at the various diffusion sensitizations, providing insight into their stability across fields of view, hemispheres, and over time. As expected, the mean SNR decreases with increasing diffusion sensitization and was estimated at $SNR_{DWI\ b=1500} = 18.5 \pm 4.8$, $SNR_{DWI\ b=4500} = 13.2 \pm 3.6$, $SNR_{DWI\ b=8000} = 10.6 \pm 2.8$ for b = 1500 $s.mm^{-2}$, b = 4500 $s.mm^{-2,}$ and b = 8000 $s.mm^{-2}$, respectively.

While SNR values remained consistent across the different modalities, including aMRI/dMRI and their different shells, intra-modality variability was observed. This variability in SNR between all FOVs for the same modality is due to the difficulty of creating homogeneous regions of the same nature close to the centers of the fields of view, due to the anatomy itself and the cutting of the blocks, thus leading to the definition of certain regions far from the center and therefore affected by the sensitivity loss of the antenna. Despite this, a consistency in SNR distributions was observed between the right and left hemispheres. This stability assesses the robustness of the scanning tissue conservation protocols over time.

dMRI acquisitions were ordered from the highest to the lowest b-value. dMRI is known to put the greatest strain on the gradient coils. A slight deviation of the SNR was systematically observed during the acquisition of the first diffusion-weighted MRI volume at b = 8000 $s.mm^{-2}$ (supplementary Fig. 7B). It is due to the application of strong diffusion gradients known to induce eddy currents within the magnet tunnel. This causes heating of the MRI passive shims, which induces a slight drift of the static field, resulting in translations of the image along the phase axis. Despite the use of a 3D EPI sequence with real-time measurement of potential field drifts, such translations could be observed during the acquisitions of the first two to three diffusion-weighted volumes at b = 8000 $s.mm^{-2}$, before stabilizing. To ensure a robust estimate of local diffusion models, the whole diffusion MRI dataset was therefore preprocessed to detect and correct such artifacts, using dedicated processing to detect outliers and correct for B0-drift-induced translations along the phase axis.

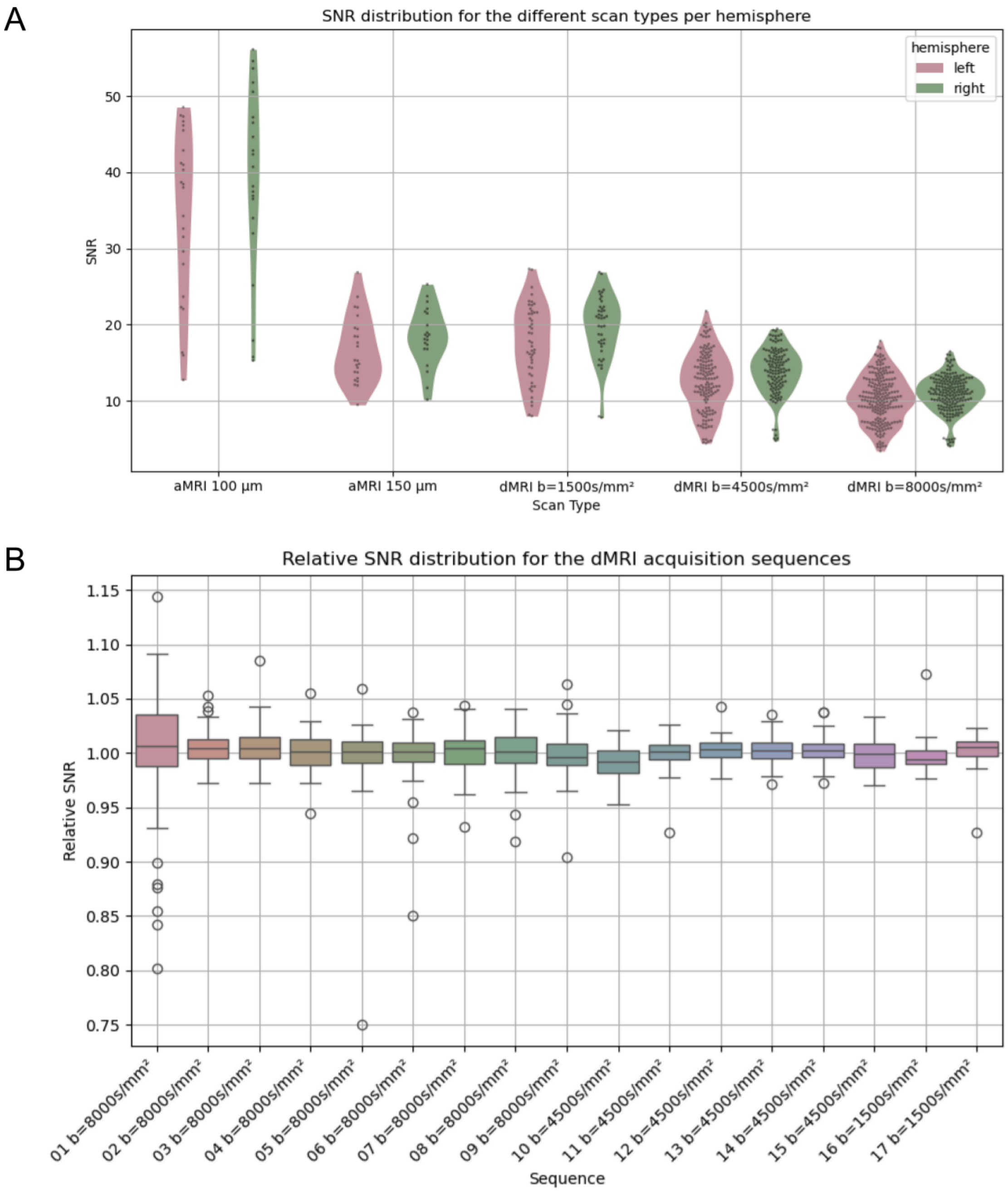


**Supplementary Fig. 7: SNR characterization of the acquisition campaigns.** Top: SNR distribution per scan type across all fields of view. Bottom: distribution of the relative SNR across all diffusion-weighted scans. Relative SNR is defined as the SNR normalized by the average SNR value within each diffusion shell per FOV, allowing comparison across FOV and across different b-values.

Supplementary text 2

**FOV Preprocessing**

*Noise filtering*

Despite obtaining acceptable SNRs for all b-values, a non-local means (NLM) filter (*74*) adapted to Rician noise was applied to enhance the quality of the diffusion MRI dataset acquired with a volume coil. This approach was preferred over alternative methods as it minimizes the number of assumptions made about the nature of the signal and can be applied equally to both quantitative and multiple-shell diffusion MRI datasets. Due to their higher SNR, anatomical maps were not filtered and left unchanged.

Parameters of the NLM filter were chosen as follows: local means were computed in blocks of size 17×17×17, considering a 8-neighborhood, with a degree of filtering set to $d_{filtering} = 0.52$, a number of antenna loops set to $n_{antenna, loops} = 1$ consistent with the volumetric nature of the antenna, and assuming a Rician model consistent with the standard reconstruction method (*e.g.*. without parallel imaging) used for the dataset. The standard deviation of the noise for each FOV was set to the value identified to compute the SNR in each FOV.

*Bias field removal*

The use of a volume coil at both 7T and 11.7T significantly reduced the B1 intensity bias, although it did not completely eliminate it at the edges of the field of view. Consequently, a bias field correction was performed on all FOVs independently using the ANTs N4BiasFieldCorrection tool (*75*) with default parameters. This correction was applied to qMRI and aMRI data. This step was essential for achieving high-quality reconstructions, as it significantly helps reduce signal jumps between adjacent FOVs during recomposition. For dMRI data, this correction was not required since the features (ODF, rotationally invariant scalar, diffusion tensor) are usually normalized voxel-wise with the T2-weighted value at $b = 0$ s.mm$^{-2}$, which is not sensitive to such intensity bias.

*Eddy currents and susceptibility artifact considerations*

The multiple-shell diffusion MRI protocol was designed using a 3D segmented echoplanar PGSE sequence with a very high number of segments (30 segments) compared to conventional imaging protocols, where the number of segments seldom exceeds 10. This designed protocol thus made the EPI data almost insensitive to any susceptibility effect or eddy current. It practically resulted in the absence of visible geometrical distortion. The price to pay for this immunity to local variations of the static field is a relatively long acquisition time (82h) to acquire the 177 diffusion-weighted volumes. The only preprocessing step required for the 177 diffusion-weighted volumes was to correct the small translation along the phase encoding direction resulting from B0 drifts induced by the heating of the passive shims present at the level of the gradient coil.

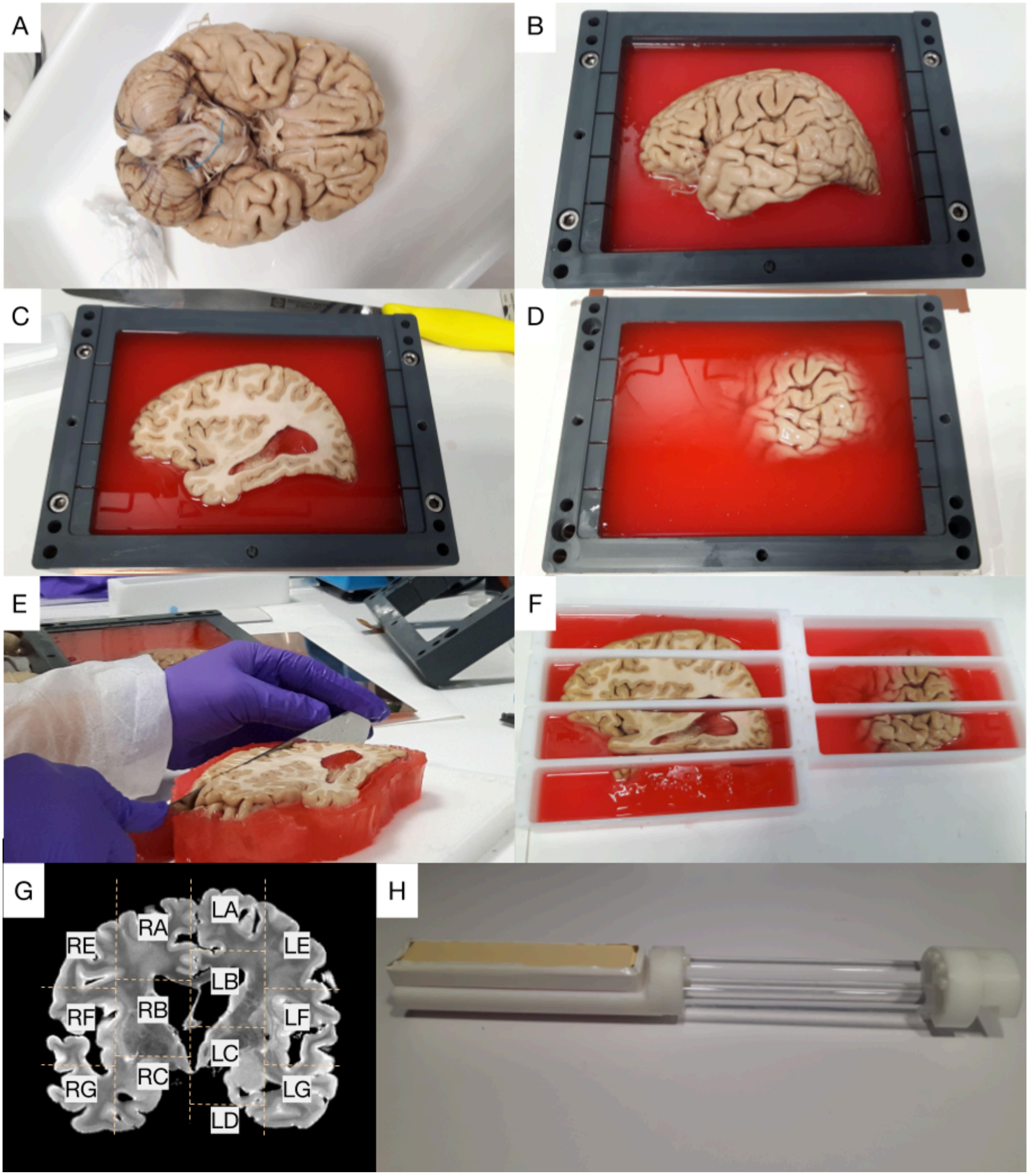


**Supplementary Fig. 8: The cutting process of the Chenonceau brain.** After fixation, the extracted 92-year-old male brain (A) was split into two hemispheres. The hemispheres were further divided into two slices of thickness 4.20cm (C, D). Six blocks were cut from the right hemisphere slices and seven from the left (E, F) and were stored in dedicated containers filled with Biz'gel. (G) represents the block nomenclature. A dedicated translating container was built to specifically fit with the 11.7 Tesla MRI and 7 Tesla MRI bores (H).

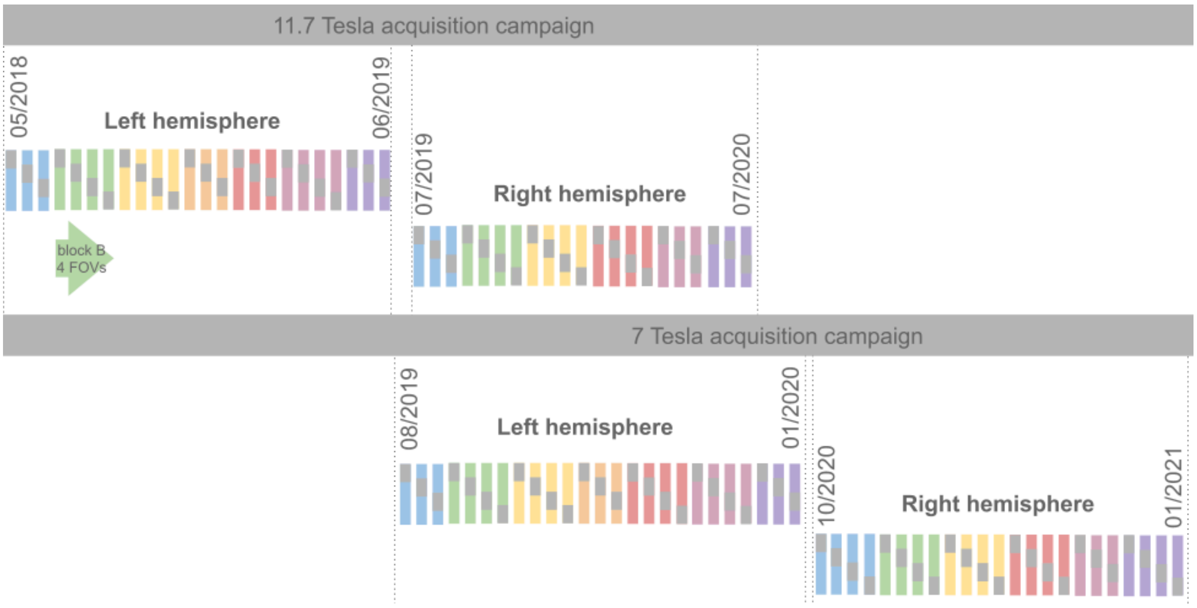


**Supplementary Fig. 9: The two Chenonceau acquisition campaigns.** The dMRI and aMRI campaign at 11.7 T accumulated over 4,800 hours of effective scanning time over a period of 28 months. In parallel, the qMRI campaign at 7 T involved more than 3,100 hours of scanning and was conducted over 18 months. Together, these campaigns constitute one of the most extensive *post-mortem* MRI acquisition efforts on the same sample.

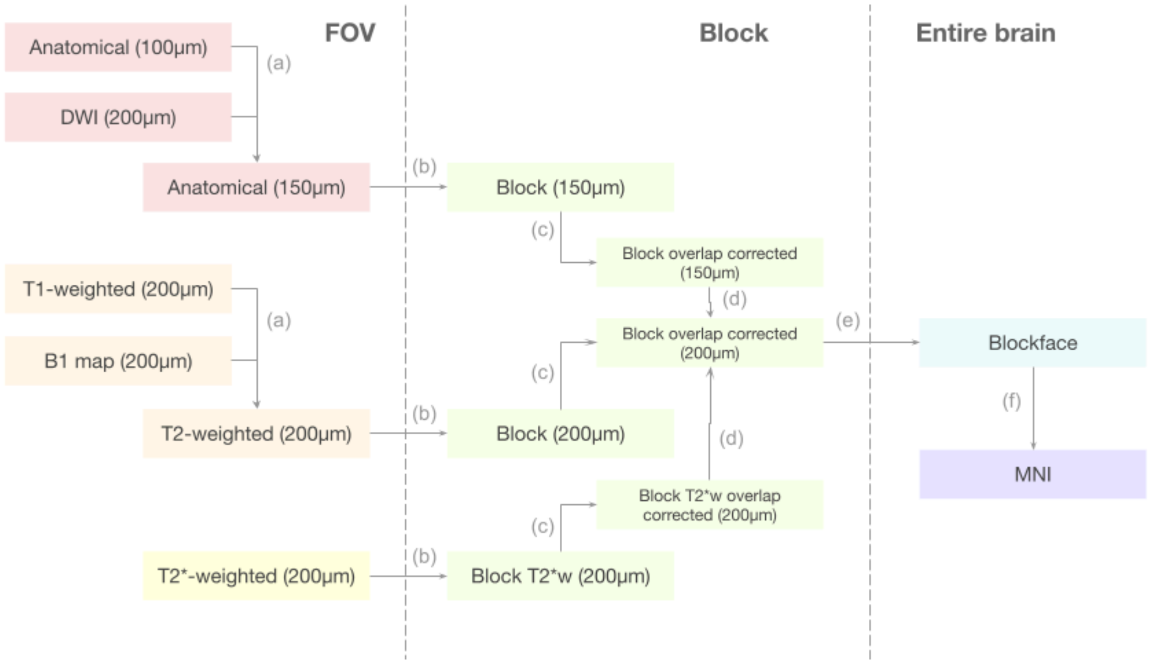


**Supplementary Fig. 10: Transformation graph from the FOVs to the blockface.** Colors represent spaces where images can be gathered. Each of the arrows represents at least one transformation: (a) the FOV-to-FOV transformation, (b-d) the FOV-to-block transformations, (e) the block-to-blockface transformation, and (f) the blockface-to-MNI transformation.

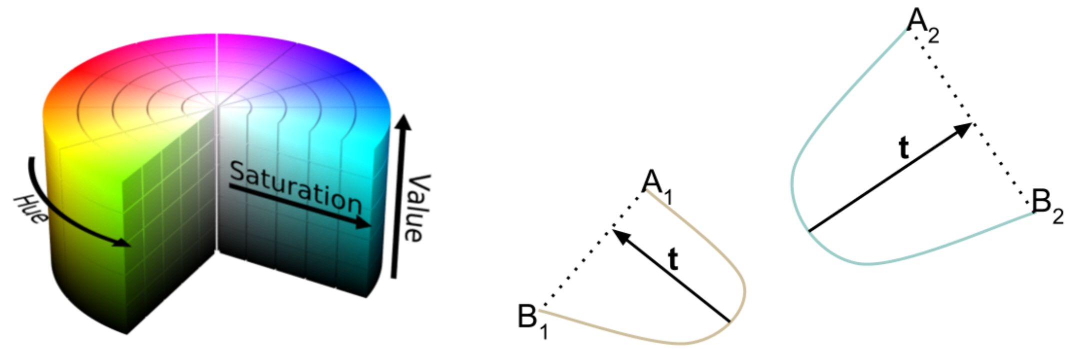


**Supplementary Fig. 11: HSV color-encoding of fiber based on tortuosity.** The tortuosity vector **t** of a fiber is defined as the vector between the furthest fiber point to the segment [AB] joining its extremities, and the corresponding orthogonal projection of this point on the segment [AB]. As an oriented vector, **t** is a good candidate for more detailed 3D color-encodings such as HSL or HSV (*71*).
